# Influence of formaldehyde on signaling pathways when used in mammalian cell culture

**DOI:** 10.1101/2024.09.17.613450

**Authors:** Katharina Ostmann, Annette Kraegeloh, Wilfried Weber

## Abstract

Formaldehyde is the smallest existing aldehyde, a highly reactive color less gas at room temperature and ubiquitously present in our atmosphere. Because of its reactivity leading to the crosslinking of macromolecules like proteins, it is widely used in industrial applications, but also in cell biology in order to preserve cells and tissues for further analysis. In this work, we show that formaldehyde releasing solutions commonly used for fixation of cells, can diffuse via the gas phase to the neighboring well and influence signaling processes in the therein cultured cells. To analyze this effect, we utilized a stable reporter cell line for YAP signaling or a gene expression-based reporter for activation of the NF-κB pathway. Especially the stable reporter cell line can also be used as sensor for bioavailable formaldehyde. The observed impact of formaldehyde on cellular signaling underscores the need for careful planning of experimental protocols and emphasizes the importance of implementing proper controls when utilizing this reagent in cellular signaling analyses.

## 1. Introduction

Aldehydes are ubiquitously present in nature, for example as aromatics in fruits or in essential oils. They are highly reactive and some are known to be carcinogenic, mutagenic and cytotoxic [1]. Chemically speaking, aldehydes are organic compounds containing a functional side group (R-CH=O) [2]. The smallest aldehyde is formaldehyde (FA). Formaldehyde is, at room temperature, a colorless gas, which is highly reactive and highly volatile, with a boiling temperature of -19 °C [3–5]. Chemically, formaldehyde reacts with macromolecules, such as proteins, in two steps. First, FA forms a covalent bond with a nucleophilic group of the molecule, leading to a methylol adduct which is converted to a Schiff base. This base can further react with another functional group of e.g. a different macromolecule, leading to a methylene bridge cross-linking the molecules [6]. FA is commonly used in a range of industrial applications including wood production or food preservation and in products like disinfectants, fabrics or cosmetics [7–9]. Formaldehyde is one of the most produced chemicals worldwide, being used to for example produce phenol or urea resins and as disinfectant and fungicide [10,11]. Ubiquitously present, atmospheric FA stems mainly from exhaust gases from cars or planes, or from heating plants or petroleum refineries. Indoors, formaldehyde often originates from cigarette smoke, resins used for plywood, disinfectants or open fireplaces [11]. In nature, FA is produced as oxidation product of terpenes or as metabolites of some bacteria [12,13]. Additionally, in humans FA is even present endogenously, also as a result of metabolism [14]. In cell biology, formaldehyde, usually released from its polymeric form PFA (paraformaldehyde) being brought into an aqueous solution, is used as a fixative in order to preserve cells and tissues for analysis via crosslinking of proteins [15]. This is leading to the death of cells and making them easy to handle for analysis like microscopy by conserving all cellular structures [16,17]. It is known to alter physiological properties of cells and tissues, like increasing the membrane permeability or as previous studies showed, (para-)formaldehyde can influence MRI behavior of tissues by changing the water diffusion constant and altering membrane physiology [18,19]. Studies even showed that using FA as fixation agent can influence liquid-liquid phase separation, either enhancing or diminishing LLPS, with samples showing droplet-like puncta in the presence of FA [20].

Since the world health organization (WHO) recognizes its risks as human carcinogen, it published a recommendation for safe exposure, stating that a concentration of 0.08 ppm should not be exceeded for more than 30 minutes [21]. This led to several studies trying to develop sensors that can accurately detect low levels of formaldehyde, preferably highly sensitive and low cost at the same time [22]. Chung et al (2013) [23] divided existing formaldehyde gas sensors in two groups, receptor-based and transducer-based sensors. Outputs can be spectrometric, colorimetric, amperometric and so on. Limitations of currently existing sensors include cross reactivity with different substances like methanol or CO_2_; their complexity, leading to the need of complicated and expensive laboratory equipment and a short lifetime since the material is usually consumed during the reaction [22,23]. Cell-based biosensors pose some advantages over traditionally used chemical sensors, including the reusability of the materials [24]. Woolston et al. (2017) [25] for example developed a formaldehyde sensor based on *E. coli* cells with either a luciferase or GFP under the control of a formaldehyde responsive promoter. They could show an operational range of their system between 1-250 µM of formaldehyde. Limitations could be that this sensor system is not feasible for higher concentrations, and due to the use of plasmids long-term experiments might be difficult [26]. In this study, we set out to utilize HeLa cells as mammalian cell line to sense formaldehyde and give an easily detectable output via fluorescence or change in absorbance.

In this study, we observe that FA activates the YAP/TAZ (Yes associated protein/Transcriptional coactivator with PDZ-binding motif) signaling pathway, which is known to be involved in a range of signaling processes, for example including inflammatory responses, biomechanical cues (mechanosignaling), ligands like growth factors or hypoxic stress [27–32]. Accordingly, we use a stable reporter cell line for hYAP1 (human YAP1) activation to sense FA. We further use an NF-κB (nuclear factor-κB)-responsive reporter to read ot the influence of FA on this pathway involved in inflammation, immunity, cell proliferation or apoptosis [33,34]. The findings of this work show that cellular reporters can be used to detect FA but also highlight that experimental protocols should carefully be designed in order to avoid inadvertent activation of mammalian signalling pathways when performing FA-based fixation of cell samples.

## 2. Materials and methods

### Development of stable cell line

The reporter cell line was constructed as described by Ostmann et al. (submitted). In brief, letiviral particles containing eGFP-hYAP1 and H2B-mTagBFP2 were produced in HEK-293T cells and subsequently, HeLa cells were transduced. The cells were sorted for eGFP and mTagBFP2 double-positive populations.

### hYAP1 activation assays

50k mL^-1^ stably transduced HeLa cells were seeded on glass slides (Carl Roth, cat no. YX03.2) in 24-well tissue culture plates and cultivated in cell culture medium (DMEM supplemented with 10% FCS, 100 U mL^−1^ penicillin and 100 µg mL^−1^ streptomycin) for 3 hours at 37 °C and 5% CO_2_. 500 µL of PBS, PFA in PBS, acetaldehyde, glutaraldehyde, isopropanol, ethanol, acetic acid, HCl, NaOH or KOH were added in indicated concentrations in the neighboring well for 1 hour. For dose dependency analysis, cells were then fixed with 4% FA for 15 minutes at RT, washed with PBS and mounted on glass slides with Mowiol 4-88. For reversibility assay, the PBS and FA was washed out with PBS and the cells were either continued to be cultured in the same medium or washed once with fresh cell culture medium and further cultivated at 37 °C and 5% CO_2_ for indicated times. Afterwards cells were fixed and mounted as before.

### Microscopy and analysis of fixed samples

Fixed samples were imaged using a Zeiss LSM 880 inverted laser scanning confocal microscope (Zeiss, Oberkochen, Germany) equipped with a 63x oil immersion (NA=1.4) objective. For analysis, Fiji was used and images were processed manually [35]. As previously described by Ostmann et al. (submitted), First, ROIs were defined for each nucleus in the BFP channel. Those ROIs were transferred to the GFP channel, the corresponding cells were then also defined as ROIs. The measured fluorescence intensity of cytosol and nucleus were then normalized to the area and subsequently divided and plotted as box plots.

### Live cell imaging

50k mL^-1^ stably transduced HeLa cells were seeded on 4-well ibidi glass bottom slides (cat. no. 80426) and cultivated in cell culture medium for 3 hours at 37 °C and 5% CO_2_. Slides were placed in a Tokai Hit incubator set at 37 °C and 5% CO_2_ for imaging. 700 µL of a 2% FA solution was added in the neighboring well at the microscope and images were taken every minute. For imaging a Zeiss LSM 880 inverted laser scanning confocal microscope (Zeiss, Oberkochen, Germany) equipped with a 40x water immersion (NA=1.2) objective was used. Analysis was performed as before using Equation 1, 2 and 3.

### NF-κB assay

For the NF-κB assay, 50k mL-1 WT HeLa cells were seeded on glass slides in 24-well tissue culture plates and cultivated in cell culture medium for 24 hours at 37 °C and 5% CO_2_. Cells were then transfected with pAF504 (3xNF-KB response elements–P_CMVmin_–SEAP) using PEI and further cultivated for 24 hours. The cell culture medium was collected (M1) and replaced with fresh cell culture medium. Then, either PBS or 2% FA was added to the neighboring well for 1 hour. For activation with TNFα, 5 ng mL-1 was added directly in the cell culture medium for 1 hour. Afterwards, the medium was collected again (M2) and replaced with fresh medium. Cells were then further incubated for 24 hours as before and subsequently the medium was collected (M3). For SEAP assay, the collected media was transferred to a 96-well round bottom plate, heated at 65 °C for 30 minutes and centrifuged for 3 minutes at 300 g. In a 96-well plate flat bottom, 100 μL 2X SEAP buffer (20 mM homoarginine, 1 mM MgCl_2_, 21% diethanolamide, pH 9.8), 80 μL of the supernatant and 20 μL pNPP solution (120 mM pNPP in 50 ml SEAP buffer) were pipetted and measured at 405 nm in a plate reader (SpectraMax iD5) for 1 hour at 37 °C, with measurements taken every minute. Of the resulting absorption curve, the linear range was taken to calculate the slope, or the change in absorption. Following formulas (**Equation 4, 5, 6, 7 and 8**) after Schlatter et al. [36] were used for the calculation of the enzymatic activity:

Lambert-Beer’s law:

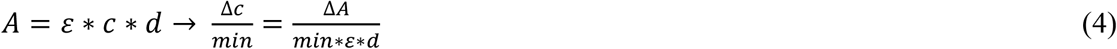

Extinction coefficient (µM^-1^ cm^-1^):

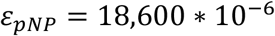

Light path (cm):

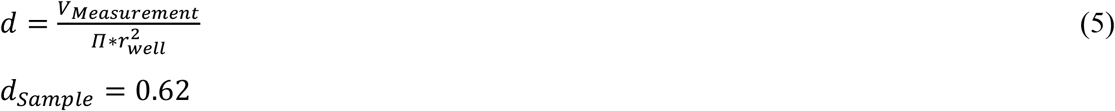

Slope (min^-1^):

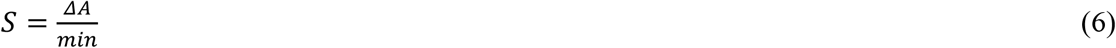

Dilution factor:

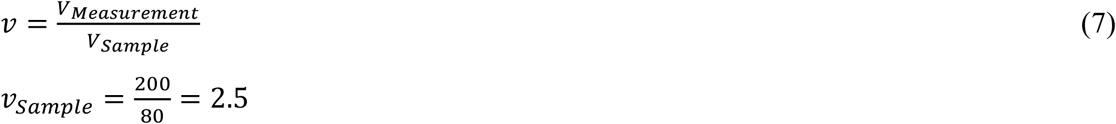

Enzymatic activity (U L^-1^, with 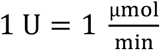):

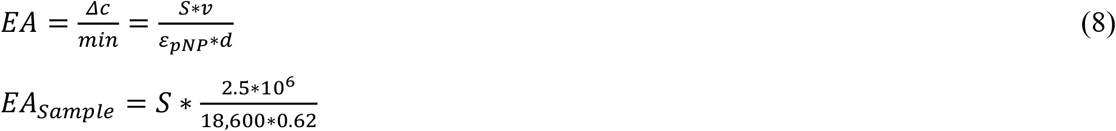

Each measurement was performed in triplicates.

### CellROX experiment

50k mL^-1^ stably transduced HeLa cells were seeded on glass slides in 24-well tissue culture plates and cultivated in cell culture medium for 2 hours at 37 °C and 5% CO_2_. The CellROX™ Deep Red Flow Cytometry Assay Kit was used and the experiment including controls performed according to the protocol provided by the manufacturer. For the samples treated with FA, FA was added to the neighboring well in indicated concentrations for 1 hour, then washed out with PBS and the staining was performed. Cells were then detached using trypsin, resuspended in FACS buffer (10% FCS in PBS) and analyzed using an Attune NxT Flow Cytometer with autosampler (Thermo Fisher Scientific). The flow cytometry results were analyzed using FlowJo™ v10.8 Software (BD Life Sciences). Cells were first gated to singlet HeLa cells, then median fluorescence as measured in the dark red channel was taken and compared between the different samples.

### Statistics

If not stated otherwise, the values indicated in the main text represent mean ± SD. For statistical analysis, single values were used and after normality tests, a Mann-Whitney U test was performed using GraphPad Prism version 9.2.0 for Windows, GraphPad Software, Boston, Massachusetts USA, www.graphpad.com.

Figure 1 was created using biorender.com.

**Figure 1.**
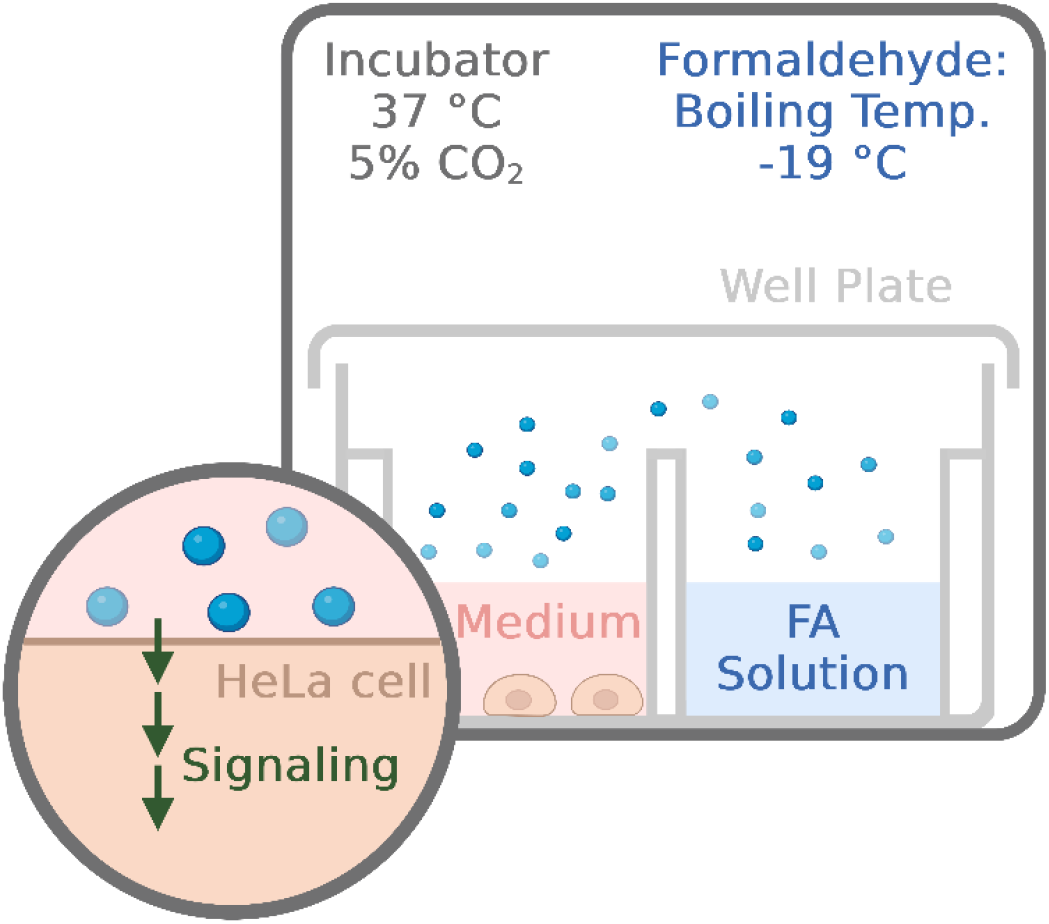
Schematic overview. Signaling in HeLa cells cultured in tissue culture well plates might be influenced by adding a formaldehyde (FA) solution to the neighboring well. FA is a colorless gas with a boiling temperature of -19 °C and is often used in cell biology as a fixative agent in order to preserve cells and tissues for analysis.

## 3. Results and Discussion

### 3.1. Low amounts of FA can activate YAP signaling

When cultivating mammalian cells in tissue culture well plates, we hypothesize that formaldehyde originating from a paraformaldehyde solution (PFA) can diffuse via the gaseous phase and activate signaling pathways in the cells cultured in the neighboring well (**Figure 1**).

First, we tested different substances and their influence on YAP signaling. For easy readout, we used a reporter cell line stably transduced with YAP1 fused to enhanced green fluorescent protein (eGFP-hYAP1) as well as histone 2B fused to the monomeric blue fluorescent protein mTagBFP2 (H2B-mTagBFP2). H2B is a histone localizing to the nucleus, hereby serving as a permanent nuclear marker, making it feasible to analyze nuclear transition of eGFP-hYAP1. We cultivated the HeLa reporter cells in 24-well plates and then added different concentrations of different aldehydes, alcohols, acids and bases to the neighboring wells. After one-hour incubation, the cells were fixed and analyzed using a confocal microscope. When using a 4% (1.33 M) and 2% (667 mM) FA solution, it can be seen that YAP localizes to the nucleus compared to the sample incubated with PBS alone (**Figure 2A**). For 4% glutaraldehyde (400 mM) or 4% acetaldehyde (909 mM) YAP also localized to the nucleus. Nuclear localization appeared to be, however not as prominent compared to when FA was used. For glutaraldehyde this could be due to a much higher boiling temperature and lower vapor pressure compared to formaldehyde, resulting in a less effective transfer (**Table 1**). Also, acetaldehyde has a higher boiling point and lower vapor pressure, although more comparable to formaldehyde. One reason for differential activation could be differences in reactivity or membrane permeability. But this still shows that other volatile aldehydes can also activate signaling in cell culture. We additionally tested different substances besides aldehydes to see if these can influence YAP (**Figure S2**). Neither 20% ethanol nor 20% isopropanol were able to activate YAP signalling, albeit being highly volatile. 10% acetic acid, 1 M HCl, 1 M NaOH and 1 M KOH did not induce nuclear localization either. This suggest an activation of YAP signaling that is specific for aldehydes.

**Table 1.** Different aldehydes and their physical properties.

| Formula | Name | Boiling Point (°C) | Vapor Pressure (kPa at 20°C) |
| --- | --- | --- | --- |
| CH<sub>2</sub>O | Formaldehyde | -19 | 430-440 [37] |
| C<sub>2</sub>H<sub>4</sub>O | Acetaldehyde | 20 | 101 [38] |
| C<sub>5</sub>H<sub>8</sub>O<sub>2</sub> | Glutaraldehyde | 187 | 2.3 [39] |

**Figure 2.**
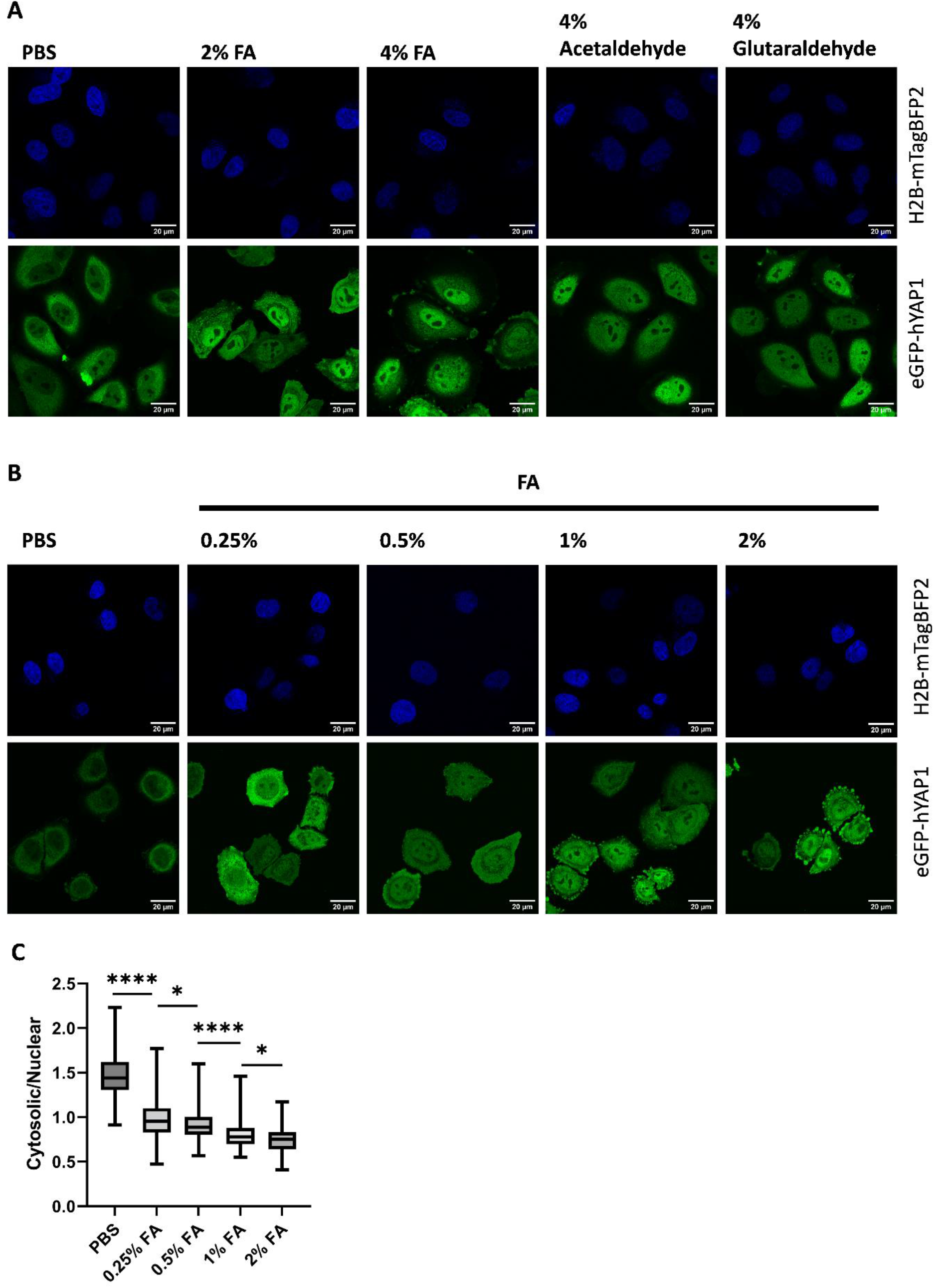
Different amounts of FA activate YAP signaling **A** Different aldehydes can activate YAP signaling in stably transduced HeLa cells with eGFP-hYAP1 and H2B-mTagBFP2. The cells were cultivated on top of glass slides for 3 hours in 24-well plates. Subsequently, PBS or aldehydes as indicated were added for 1 hour to the neighboring well prior to fixation of cells and analysis by confocal microscopy. Scale bar: 20 µm. **B** Different concentrations of FA (diluted in PBS) were used to test for activated YAP signaling. The experiment was performed as before mentioned. Scale bar: 20 µm. **C** Quantitative analysis of nuclear localization of YAP. The ratio of fluorescence intensity of nuclear to cytosolic YAP was determined by image analysis. Data in C are representative data from n = 3 replicates. Experiments were performed in duplicates and at least four randomly distributed images were taken from each sample. Boxes represent median and upper and lower quartiles, whiskers represent min and max values. Mann-Whitney U test, \*\*\*\**P* ≤ 0.0001, \**P* ≤ 0.05

Next, we wanted to investigate which concentrations of FA can influence YAP signaling. All used concentrations of FA (0.25% - 4%) were sufficient to activate YAP signaling, seen by nuclear localization of hYAP1 (**Figure 2B**). At higher concentrations, the cells form vesicles and excesses, which are also visible at lower concentrations but much less prominent. Some vesicles also formed in the PBS control, but here, hYAP1 did not localize to the nucleus. The images were then further processed by quantifying the cytosolic/nuclear ratio of YAP localization (**Figure 2C**). The differences of localization between the used FA concentration are statistically significant (*P* ≤ 0.05), suggesting a high sensitivity, with low amounts being sufficient to activate signaling and additionally small differences in concentrations can be distinguished. Since this reaction is presumably activated by diffusion of FA via the gaseous phase to the next well, it can be assumed that the actual concentration in the cell culture medium is much lower than what is originally present in the used solutions.

### 3.2. Live Cell Analysis of FA-induced signalling

To characterize the time-course of activated YAP signaling using FA, live cell imaging was performed. To this end, the reporter cells were cultivated, 2% FA solution was added to the neighboring well and the cells were imaged directly. It can be observed, that localization to the nucleus starts after approximately 20 minutes (**Figure 3A and Figure 3B**). After around 50 minutes, nuclear localization reached a plateau. Also, the cells seem to lose cell-cell contacts and start to form vesicles and excesses as seen before, maybe suggesting membrane blebbing due to ongoing apoptosis [40]. Cell shrinkage has also been described as an early onset apoptotic marker, leading to the separation of the dying cell from the surrounding cells, also supporting the theory of ongoing apoptosis [41]. It is however also described that in cells undergoing apoptosis and membrane blebbing, nuclear fragmentation occurs at the same time or even before blebbing. This cannot be observed in our cells, which could either be due to the fact that 1 hour is not enough for nuclear disintegration or that another mechanism is at play. Starting necrosis could also lead to membrane blebbing, but in that case, the membrane becomes leaky and intracellular molecules get released, which at least for our reporter molecule couldn’t be observed [42].

**Figure 3.**
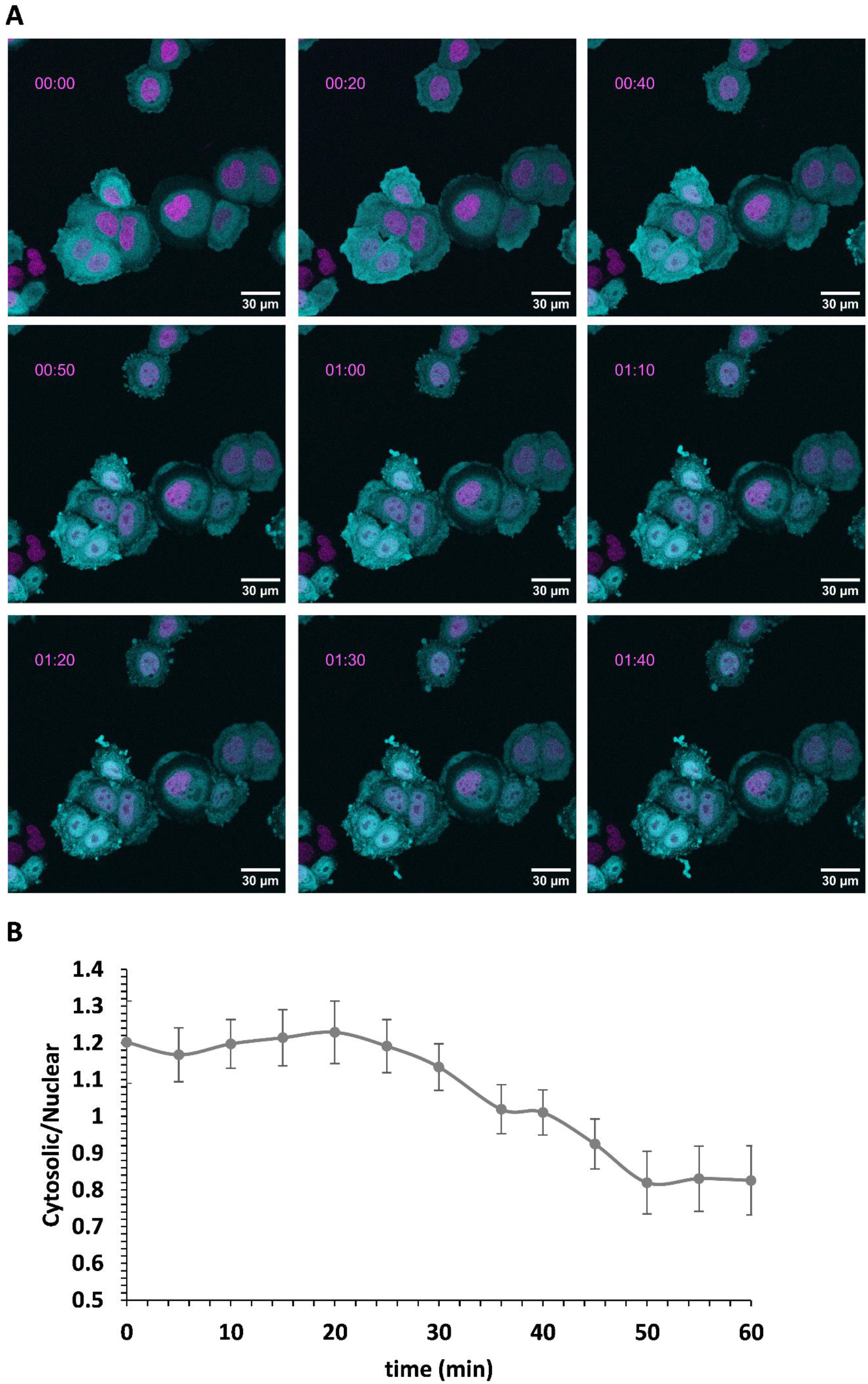
Live cell imaging of FA activating YAP signaling **A** The stably transduced HeLa cells (eGFP-hYAP1 and H2B-mTagBFP2) were seeded in 4-well ibidi slides and cultured for 3 hours. 2% FA was then added directly at the microscope and an image was taken every minute. Time stamp: hh:mm. Scale bar: 30 µm. **B** Quantitative analysis of nuclear localization as before. Data points represent the mean ± s.d.

### 3.3. Activated Yap signaling through FA is reversible

A study that could support the theory that the cells undergo apoptosis, showed that ethanol induced apoptosis is reversible after 2 hours when washed and incubated in fresh medium [43]. Consequently, the next aim was to examine if FA induced membrane blebbing and additionally nuclear YAP localization in our reporter cells can also be reversed. In order to test for the reversibility of this reporter system, cells were cultivated as before and 2% FA solution was added in the proximate well. After 1 hour, FA was washed out with PBS, the cells were washed with medium once and then further cultivated for 2 hours to 48 hours. **Figure 4** shows that already 4 hours after washing, hYAP1 localized in the cytosol again. It could also be observed, that membrane blebbing almost completely diminished after 4 hours. This could, as mentioned before, indicate the start of apoptosis in FA treated cells which is reversed after washing.

**Figure 4.**
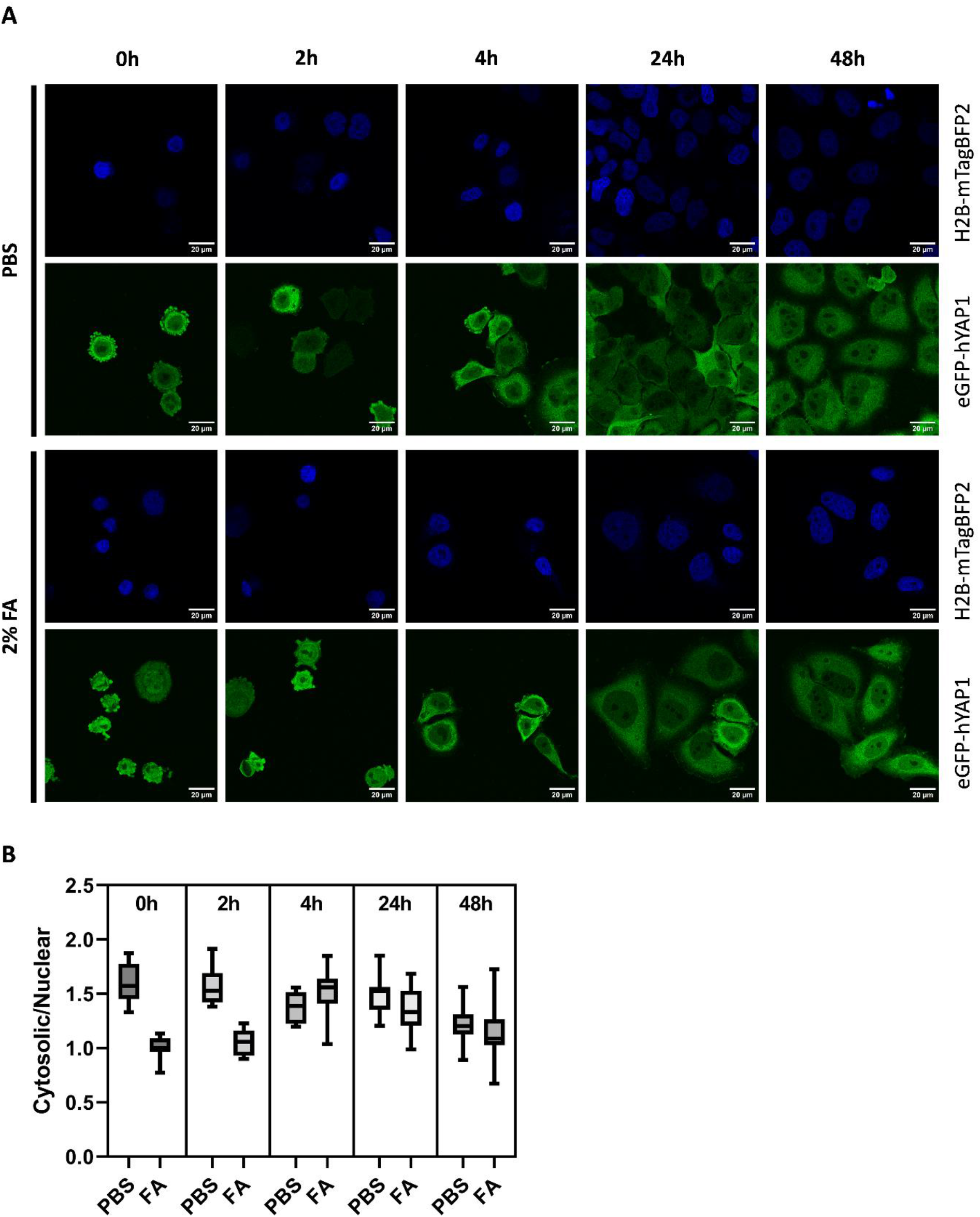
Reversibility assay of YAP signaling after PFA activation. **A** HeLa cells stably expressing eGFP-hYAP1 and H2B-mTagBFP were cultivated on top of glass slides for 3 hours. Subsequently PBS or 2% FA was added for 1 hour to the neighboring well. First samples were taken at this point labelled “0 h”. The FA was washed out with PBS and the medium was taken off, the cells were washed once with medium and cultivated in fresh medium for different time spans (2h – 48h). Scale bar: 20 µm. **B** Quantitative analysis of nuclear localization as before. Data are taken from n = 3 different image sections. Boxes represent median and upper and lower quartiles, whiskers represent min and max values.

Interestingly, after removal of the FA solution from the neighboring well but without exchange of the medium covering the cells, no reversibility could be observed even after 48 hours (**Figure S1**). This suggests that the medium itself contained either FA or some reaction products which were still active after at least two days.

### 3.4. FA also activates the NF-κB pathway

To further investigate whether FA also activates different signaling pathways, we used a reporter previously developed by our group [44]. This reporter is dependent on NF-κB signaling, where an activation of the pathway leads to the expression of secreted alkaline phosphatase (SEAP). In transiently transfected HeLa cells, FA led to a significantly higher expression of SEAP compared to the negative control using PBS (**Figure 5**) and reaching 70% of activation by tumor necrosis factor alpha (TNFα) [45].

**Figure 5.**
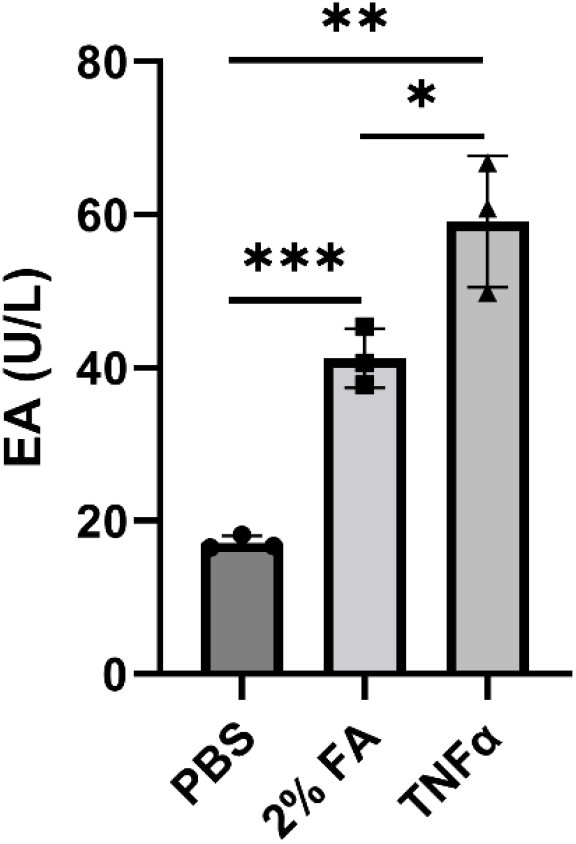
NF-κB activation through FA. HeLa cells were transiently transfected with an NF-κB responsive expression vector for secreted alkaline phosphatase (SEAP). PBS or 2% FA were added to the neighboring well for 1 hour. As control, 5 ng mL^-1^ TNFα (tumor necrosis factor α) was added directly to the medium for 1 hour. Afterwards, the medium was taken off and replaced with fresh medium, respectively. PBS and FA were washed out of the well. After 24 hours, the supernatant was taken for analysis and a SEAP assay was performed.

As to why signaling gets activated through FA, we hypothesize different possibilities. First, since FA is a crosslinking agent, it could be that extracellular structures like growth factors or receptors are crosslinked and thus activate signaling [17]. In the case of YAP signaling it could also be that, through crosslinking, the cell structures become stiffer, leading to activation through mechanosignaling. A study by Kim et al. (2017) [46] investigated if the stiffness of cells increases upon direct treatment with FA. They could show that concentrations of FA between 0.1 to 10% led to an increase of the measured Young’s modulus of the individual cells, while concentrations below 0.0001% FA did not lead to an increase in stiffness. Since Kim et al. did not use concentrations of FA between 0.0001% and 0.1% it is difficult to say whether concentrations in between actually lead to stiffening of cells, but it could still be an explanation why YAP signaling gets activated.

Studies also showed that aldehydes lead to an increase in reactive oxygen species (ROS), leading to oxidative stress responses, including activation of YAP and NF-κB signaling. Usually, aldehyde dehydrogenase 2 (ALDH2) catalyzes the reaction from toxic formaldehyde to non-toxic products [47]. In HeLa cells, it is known that the expression levels of ALDH2 are low [48]. This could lead to a build-up of ROS once aldehyde is present in the medium and in the cells and thus lead to an increased YAP and NF-κB signaling. Since oxidative stress is also a driver of apoptosis, it could explain the morphology of the FA treated cells [49]. However, using a CellROX kit, which can indicate ROS, we could not show a production of ROS when treated with FA (**Figure S5**). This theory still seems to be the most likely scenario and should not yet be disregarded, since ROS formation is dependent on a lot of factors, ROS levels are often very low and they usually have a short half-life making them difficult to proof [50].

## 4. Conclusion

Overall, our findings suggest that FA can activate YAP and NF-κB signaling This reaction was fast, sensitive and could be reversed, making the biosensor easy to handle and reusable.

Further investigations will be made in order to fully comprehend the influence formaldehyde has on cells and/or signaling processes. As a side note, with our findings in mind, we recommend the use of FA as fixation agent only with a spatial distance between all samples, especially if they are only partially fixed while other samples remain in culture.

## Acknowledgments

We thank the staff of the Life Imaging Center (LIC) in the Center for Biological Systems Analysis (ZBSA) of the University of Freiburg for help with microscopy resources, and the excellent support in image recording and analysis. We acknowledge the excellent scientific and technical assistance of the Signalling Factory Core Facility staff of the Albert-Ludwigs-University Freiburg for help with the plate reader and flow cytometry. We thank the Lighthouse Core Facility for their support with cell sorting. LCF is funded in part by the Medical Faculty, University of Freiburg (Project Numbers 2021/A2-Fol; 2021/B3-Fol) and the DFG (Project Number 450392965).

## Author contributions:CRediT

**Katharina Ostmann**:Writing – original draft, Methodology, Investigation, Formal analysis, Data curation. **Annette Kraegeloh**: writing – review and editing, Validation, Supervision. **Wilfried Weber**: writing – review and editing, Validation, Funding acquisition, Supervision, Project administration, Resources, Investigation, Data curation.

