## Supplementary-Information for "Influence of formaldehyde on signaling pathways when used in mammalian cell culture"

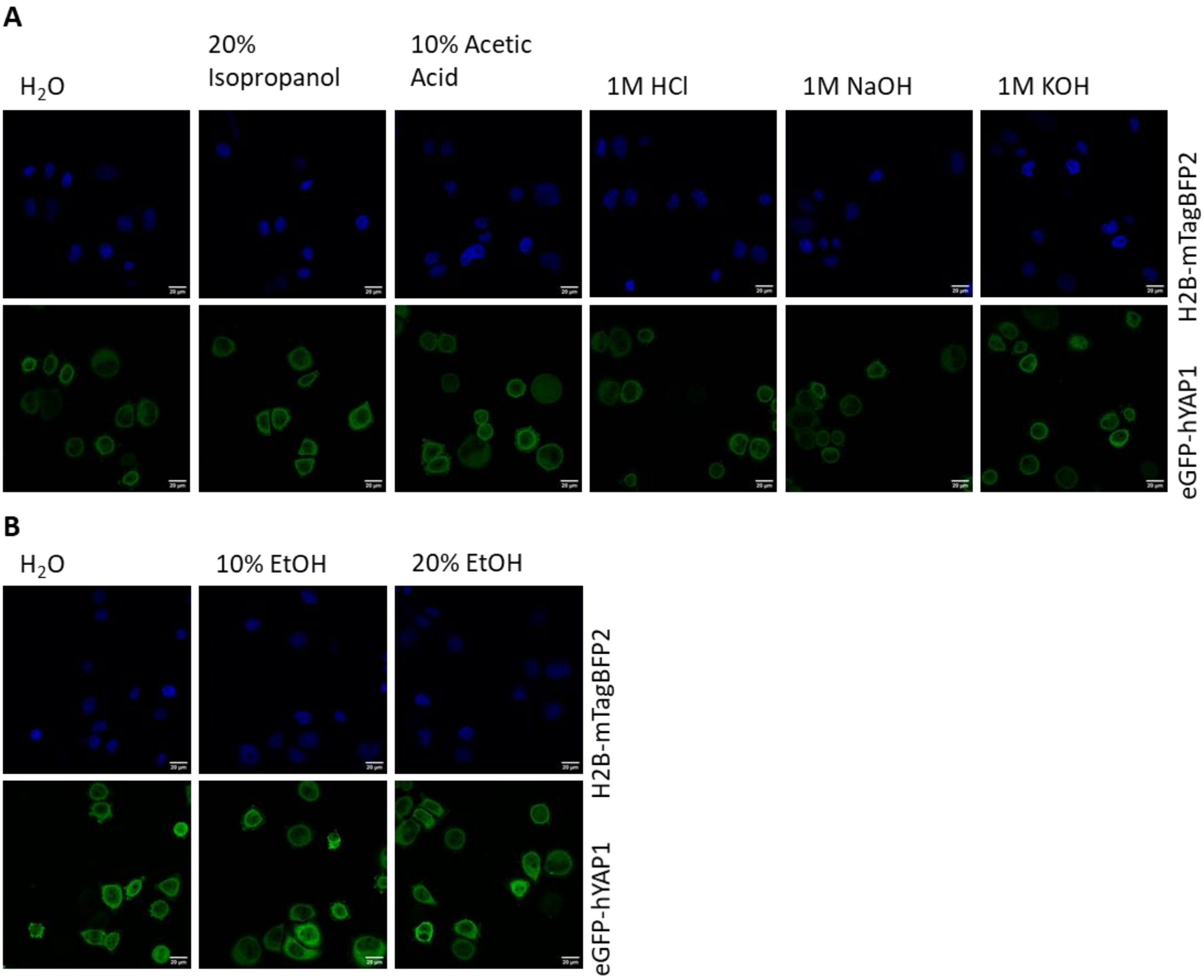


**Figure S1.** Activation of YAP signaling by different substances. The indicated substances were tested whether they can activate YAP signaling in stably transduced HeLa cells with eGFP-hYAP1 and H2B-mTagBFP2. The cells were cultivated on top of glass slides for 3 hours. Subsequently, PBS (negative) or indicated substances in indicated concentrations were added for 1 hour to the neighboring well, respectively. Please note that experiments were performed on different days. Scale bar: 20 µm.

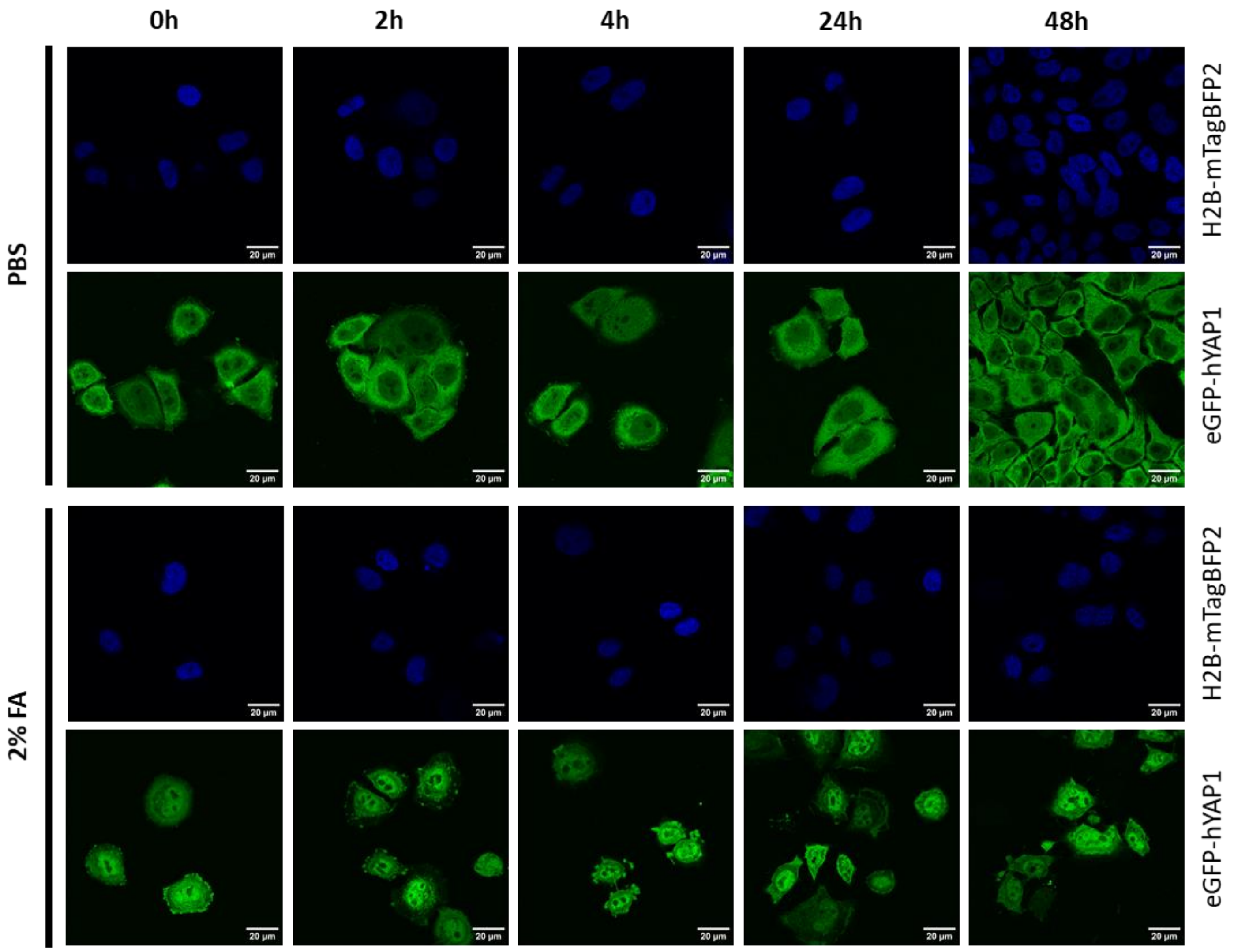


**Figure S2.** Reversibility of FA activated YAP signaling without washing. HeLa cells stably expressing eGFP-hYAP1 and H2B-mTagBFP were cultivated on top of glass slides for 3 hours. Subsequently PBS or 2% FA was added for 1 hour to the neighboring well, respectively. First samples were taken at this point labelled “0 h”. FA was washed out with PBS and the cells were cultivated in the same medium for different time spans (2h – 48h). Scale bar: 20 µm.

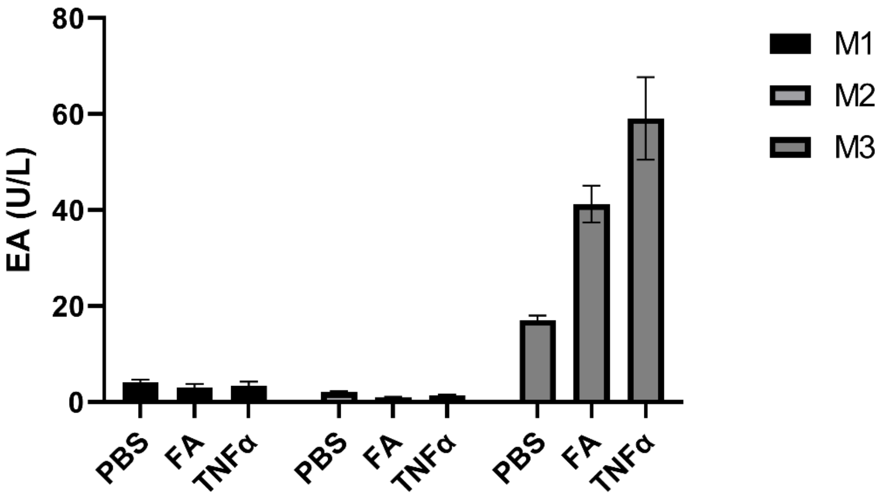


**Figure S3.** Influence of FA on NF-κB signalling. HeLa cells were transiently transfected with the NF-κB responsive SEAP reporter. The medium was collected (M1) and replaced with fresh medium. PBS or 2% FA were added to the neighboring well for 1 hour. As control, TNFα (tumor necrosis factor α) was added directly to the medium for 1 hour. Afterwards, the medium was taken off (M2) and replaced with fresh medium, respectively. PBS and FA were washed out of the well. After 24 hours, the supernatant (M3) was taken for analysis and a SEAP assay was performed.

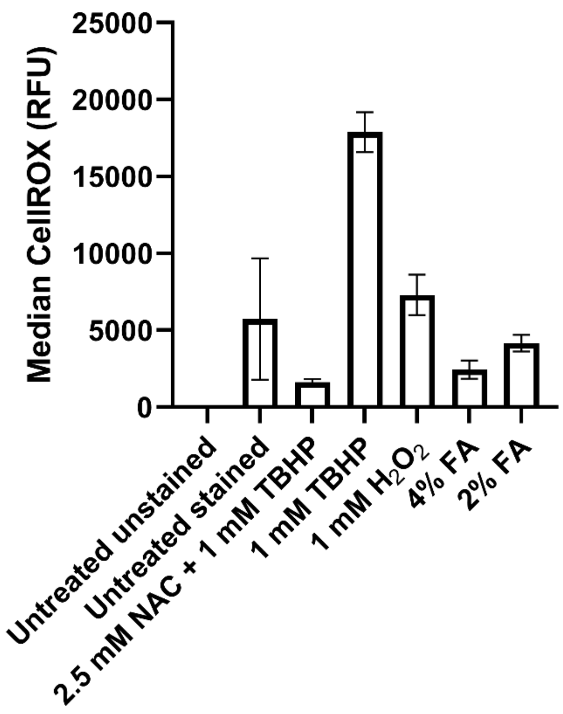


**Figure S4.** CellROX assay to investigate ROS formation. CellROX Deep Red staining was performed using the HeLa reporter cell line. As negative controls, untreated cells were analyzed and additionally NAC (N-acetyl cysteine) was used as an antioxidant. Positive controls were TBHP (tert-butyl hydroperoxide) and $H_2O_2$, known to induce ROS formation in cells. Following 1 hour incubation with possible ROS inducers, cells were stained and subsequently analyzed using flow cytometry. Cells were gated to single cells and the median CellROX Deep Red fluorescence was measured.
